# Evaluation of Neuronal Activation Thresholds for Low-Frequency Electromagnetic Exposure Using Morphologically Realistic Neuron Models

**DOI:** 10.64898/2026.04.17.719188

**Authors:** Joaquín Gázquez, Carolina Camacho Cadena, Wenzhe He, Eikei Yamada, Carsten Alteköster, Florian Soyka, Ilkka Laakso, Akimasa Hirata, Wout Joseph, Thomas Tarnaud, Emmeric Tanghe

## Abstract

International guidelines for low-frequency electromagnetic field exposure (LF EMF) are primarily intended to prevent substantiated adverse effects. In the frameworks, limits on internal electric fields are linked to external exposure levels through computational dosimetry. However, the relationship between internal electric fields and these adverse effects remains incompletely understood. In particular, current approaches often overlook the morphological complexity and diversity of cortical neurons, which may limit the realism of neuronal activation estimates used to support these assessments. This study evaluates LF EMF-induced neural activation using 25 morphologically realistic neuron models spanning all cortical layers, embedded within 11 detailed human head models. The internal electric fields were simulated for uniform magnetic field exposures (100 Hz–100 kHz) along the three anatomical directions, and excitation thresholds were computed using a multi-scale framework combining voxel-based dosimetry with biophysical neuron simulations. A real-world exposure scenario involving a child near an acousto-magnetic article-surveillance deactivator was also analyzed. Thresholds varied across cell type, morphology, cortical location, subject anatomy, frequency, and exposure direction, with L2/3 pyramidal, L4 basket, and L5 thick-tufted pyramidal cells showing the lowest thresholds. Despite this variability, all simulated thresholds were conservative with respect to the basic restrictions and dosimetric reference limits set by IEEE ICES and ICNIRP. The smallest margin occurred at 100 kHz, where the threshold remained a factor of 2.8 above the corresponding limit. These findings indicate that current LF EMF exposure limits remain conservative when evaluated using highly detailed, morphology-based CNS activation models.

## I. Introduction

TO protect people from adverse health effects, the IEEE International Committee on Electromagnetic Safety (ICES) and the International Commission on Non-Ionizing Radiation Protection (ICNIRP) have established international standards and guidelines that limit the permissible exposure to electromagnetic fields (EMF) across a wide range of frequencies [1], [2]. For human exposure to EMF with frequencies up to 100 kHz, unintended neuromodulation of the central nervous system (CNS) and pain in the peripheral nervous system (PNS) are regarded as substantiated adverse health effects with the lowest threshold [1], [2].

In this study, low frequencies (LF) are defined as EMFs in the range of 100 Hz to 100 kHz, which are commonly used in applications such as induction heating, wireless power transfer (WPT) and electronic article surveillance (EAS) systems [3]– [8]. Exposure to LF EMFs induces electric fields in living tissues, which may modulate neural activity in both the CNS and PNS. Depending on factors such as the location, duration, and type of neural activation, such unintended neuromodulation may lead to adverse health effects [9]–[11].

In international exposure guidelines, the assessment process first identifies the thresholds for adverse health effects. After applying reduction (safety) factors, limits on internal physical quantities, most notably the internal (*in situ*) electric field for LF EMF, are established as basic restrictions (BRs) by the ICNIRP or dosimetric reference limits (DRLs) by the IEEE ICES. These internal limits are then used to derive the permissible exposure levels in terms of external electric and magnetic field strengths, referred to as reference levels (RLs) by the ICNIRP or exposure reference levels (ERL) by the IEEE ICES [1], [2].

The derivation of exposure limits from internal quantities relies on realistic numerical simulations of human exposure to external electric and magnetic fields. This process involves anatomically detailed head models assigned with the dielectric properties of head tissues, numerical methods such as the scalar potential finite difference (SPFD) method and the finite element method (FEM) to compute the internal electric field induced by external fields, and activation models to evaluate neuronal firing in response to the induced electric field [12]– [14]. The smallest internal electric field capable of eliciting neural activation is known as the excitation threshold. Although neural activation does not necessarily constitute an adverse health effect, it serves as an important physiological reference for understanding the mechanisms underlying the perception, discomfort, and pain induced by low-frequency EMF exposure. Accordingly, the excitation thresholds derived from neural models are used as supporting evidence in the derivation of exposure limits, rather than as direct criteria for guideline formulation. Exposure limits may further differ depending on the population and environmental conditions, such as restricted or unrestricted environments (IEEE ICES) and general public or occupational exposure settings (ICNIRP) [1], [2].

Exposure limits are periodically updated to incorporate the latest scientific evidence on EMF protection. At present, spatially extended nonlinear node (SENN) models are widely used in nerve activation simulations to support the scientific rationale underlying the derivation of BRs and DRLs [2], [15]. Various extensions and modifications of the SENN model have been investigated to account for the key parameters influencing axonal electrostimulation in biological neurons. The necessity for such efforts has been mentioned in a research agenda by IEEE ICES and a gaps of knowledge document by ICNIRP [16], [17].

For example, Gomez-Tames et al. demonstrated differences in excitation thresholds across multiple membrane models and quantified the impact of intersubject variability based on three human head models [13]. Similarly, Soyka et al. examined alternative exposure limit estimates by comparing action potential thresholds derived from five ion channel dynamics within the SENN framework, as well as from two double-cable models (MRG-Sensory and MRG-Motor) at temperatures of 22 ^◦^C and 37 ^◦^C [18], [19].

Furthermore, Gomez-Tames et al. and Soldati et al. employed multiscale computational approaches to investigate excitation thresholds in the CNS by embedding axon models into anatomically realistic phantoms [12], [20]. An overview of these extensive efforts to characterize nerve activation mechanisms under EMF exposure is provided in a review paper, which summarizes discussions from a Computational Bioelectromagnetics workshop organized by the IEEE ICES [14].

However, axon models are not representative of the full reality of CNS electrostimulation, as they do not incorporate the various morphological components of a neuron, such as the soma, dendrites, and axon collaterals. In particular, the various terminations of the axon collaterals and the bends of the corticofugal axons are highly excitable to electrostimulation [21], [22]. Furthermore, apart from variations in membrane dynamics, the morphology of neurons varies across the six cortical layers. This factor has not yet been systematically explored in the context of EMF exposure. Hence, a task group was established and led by the authors within IEEE ICES Technical committee 95 Subcommittee 6 to explore the excitability of morphologically realistic cell models in the context of electromagnetic safety in the LF range.

This study improves the assessment of physiological basis of internal fields for LF EMF exposure of the CNS by evaluating activation thresholds of morphologically realistic neuron models of multiple cortical layers incorporated into various head models. We approach this by introducing 4 key novelties into our study:

- A uniform magnetic field aligned with the 3 anatomical directions is simulated in 11 human head models to account for inter-subject variability, providing a more representative analysis than previous research [13].
- Morphologically realistic neuron models are used to evaluate a real-life exposure scenario. The induced electric field inside the head of a child standing in front of a cash register is simulated while the child is exposed to the magnetic field of a deactivator from an acousto-magnetic (AM) EAS system.
- Following the methodology of Aberra et al. (2018, 2020), in which morphologically realistic neuron models are used to investigate transcranial magnetic and microelectrode stimulation, we use 25 neuron models representing 5 cell types across 6 cortical layers, offering the most in-depth analysis of LF EMF exposure and safety assessment to date [21], [23].

Our findings contribute to ensuring that exposure guidelines are based on realistic, research-driven models that fully encompass the heterogeneity of the CNS.

### II. Methodology

#### A. Electromagnetic field simulations

##### 1) Head models

MRI-based head models of 11 healthy subjects were used in the uniform magnetic field simulations (5 female, 6 male, mean age 29.2 ± 3.1, age range 25 - 37). T1- and T2-weighted Magnetic Resonance (MR) images were acquired using a 3 T MRI scanner (Magnetom Skyra; Siemens, Ltd, Erlangen, Germany) with parameters as follows - T1- weighted: repetition time (TR)/echo time (TE)/inversion time (TI)/flip angle (FA)/field of view (FOV)/voxel size/slice number = 1800 ms/1.99 ms/800 ms/9°/256 mm/1×1×1 mm/176; T2-weighted: TR/TE/FOV/voxel size/slice number = 3200 ms/ 412 ms/256 mm/1×1×1 mm/176. For the generation of the head models, a semi-automatic segmentation pipeline was used to segment the MR images into different tissues [26]. In brief, non-brain tissues were segmented based on T1- and T2-weighted MR images, while brains were reconstructed from T1-weighted MR images using FreeSurfer image analysis software [27], [28].

##### 2) Volume Conductor Solver

For the uniform magnetic field scenario, volume conductor models were generated from segmented images by voxelizing them into 0.5 × 0.5 x 0.5 mm voxels. To account for the frequency-dependence of electrical tissue conductivities, frequencies of 300 Hz and 100 kHz were used for the electric field simulations. For frequencies between 100 Hz and 10 kHz, the electrical properties corresponding to 300 Hz were applied, whereas for frequencies between 10 kHz and 100 kHz, the properties corresponding to 100 kHz were used. At 300 Hz, the electrical conductivity values *σ* were assigned as follows with unit of S/m: grey matter (0.1), white matter (0.06), cerebrospinal fluid (2.0), cortical and cancellous bone (0.02 and 0.07), fat (0.04), muscle (0.35), dura (0.5), and blood (0.7) [29]. At 100 kHz, the electrical conductivity values were obtained from the four Cole–Cole model [30] and were (S/m): grey matter (0.1337), white matter (0.0818), cerebrospinal fluid (2.0), cortical and cancellous bone (0.0208 and 0.0839), fat (0.0484), muscle (0.3618), dura (0.5019), and blood (0.7029). The conductivities were assumed to be isotropic for both conditions.

A sinusoidal, time-varying uniform magnetic field of 1 mT along the three orthogonal anatomical orientations in both directions is applied to the head model. The electric field **E** is computed by evaluating the magnetic vector potential **A** and the electric scalar potential *ϕ*:

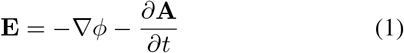

The electric potential is derived by solving the Ohmic quasistatic equation:

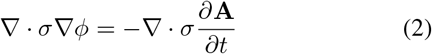

The Ohmic quasi-static formulation effectively decouples the electric and magnetic fields. This assumption holds at intermediate frequencies, where the displacement current is significantly smaller than the conduction current [31]–[33].

The unknown electric scalar potential was determined by numerically solving Equation (2) using the finite-element method with a geometric multigrid solver [34]. The elements were first order and corresponded to the 0.5 × 0.5 × 0.5 mm voxels of the volume conductor models. In solving Equation (2), the boundary condition was that the normal component of the current density was zero on the surface of the head model. The induced electric field scales linearly with frequency in a uniform time-varying magnetic field. Therefore, once the electric field was determined for a given frequency, the field at any other frequency was obtained by scaling the field proportionally to the frequency.

In this study, FEM was implemented on a structured voxel grid rather than on an unstructured tetrahedral mesh, allowing a direct comparison with the scalar potential finite-difference (SPFD) formulation. The unknown electric scalar potential was independently computed using the SPFD method [35]. In this method, the scalar potential was defined at voxel vertices, and Equation (2) was discretized by enforcing current conservation for branch currents flowing along voxel edges. The resulting linear system was solved using a solver adapted for the SPFD formulation, based on the geometric multigrid and successive over-relaxation (SOR) framework originally proposed for finite-element analyses [34]. The same boundary conditions as in the FEM analysis were applied. Figure 1(a-c) shows slice examples of the electric field using the FEM-solver.

**Fig. 1.**
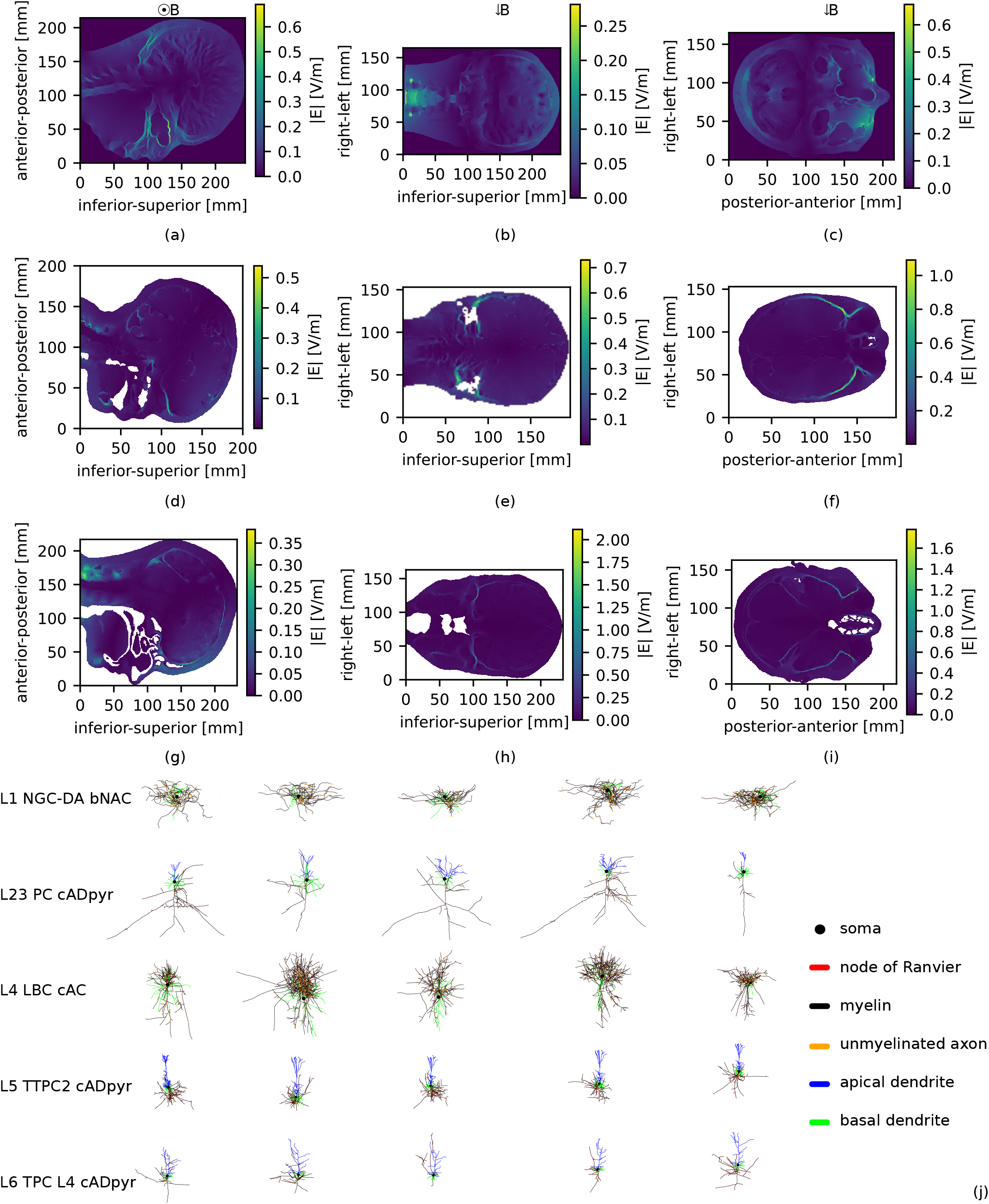
Magnitude of the electric field for a uniform magnetic field of 1 mT in the lateral-medial direction (arrow shows direction of the magnetic field) using a frequency of 300 Hz. (a) Sagittal slice. (b) Coronal slice. (c) Transverse slice. Magnitude of the electric field for an AM-EAS tag deactivation pulse using the MIDA model [24]. (d) Sagittal slice. (e) Coronal slice. (f) Transverse slice. Magnitude of the electric field for an AM-EAS tag deactivation pulse using the MARTIN model [25]. (g) Sagittal slice. (h) Coronal slice. (i) Transverse slice. (j) 2D morphological projection of the 25 neuron models. Each row corresponds to a cell type, with abbreviations indicating cortical layer, morphological class and electrical type. Five clones are used per cell type to account for morphological variability. **Abbreviations**. NGC-DA: neurogliaform cells with a dense axonal arbor, PC: pyramidal cell, LBC: large basket cell, TTPC: thick-tufted pyramidal cell with an early bifurcating apical tuft, TPC L4: tufted pyramidal cell with its dendritic tuft terminating in L4. bNAC: burst non-accommodating, cADpyr: continuous adapting pyramidal, cAC: continuous accommodating [21], [23].

##### 3) Electromagnetic field simulations for acoustomagnetic electronic article surveillance deactivator

AM-EAS deactivator systems use a short, high intensity, magnetic pulse to deactivate tags. To generate such a pulse, a coil is installed in the cashier’s desk. Therefore, a child’s head can be just at the height of the coil and be strongly exposed to the magnetic field. The dimensions used for the simulation setup are shown in Fig. 2(a,c) and were measured at a cash register in a retail shop. They therefore represent realistic worst-case conditions. The current flowing through the simulated coil was calibrated such that the simulated magnetic field matches magnetic field measurements of the real device. Furthermore, the time course of the magnetic field was measured. An exponentially damped sine wave in Fig. 2(b) was fitted to the data and used in the calculations to provide a realistic simulation setup:

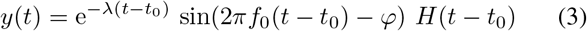

**Fig. 2.**
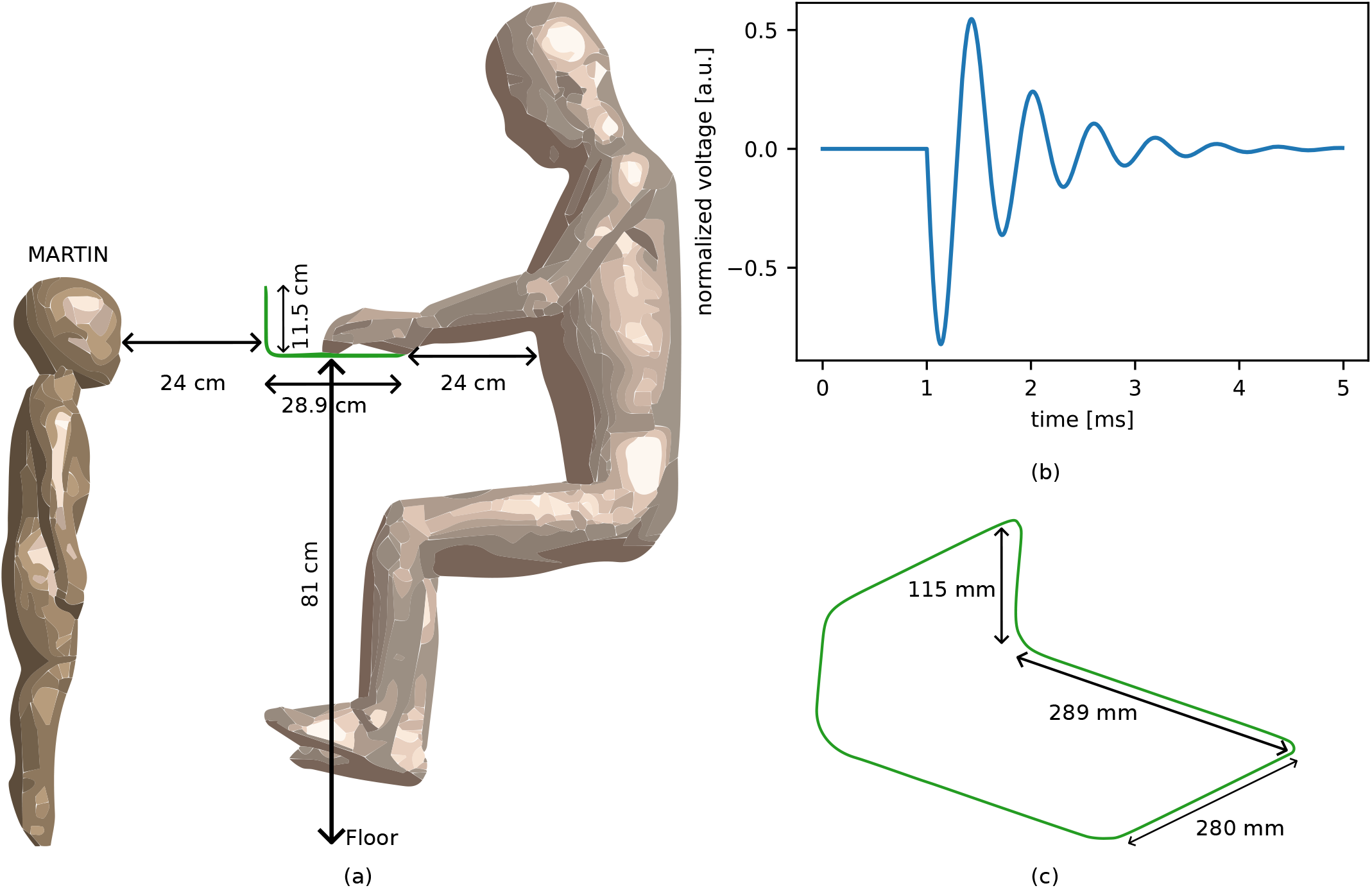
(a) Schematic reconstruction of the simulation setup used in Sim4life of a 29-month-old male toddler standing in front of a cash register during AM-EAS tag deactivation. The person on the right side is not being simulated and serves only as a reference. (b) Magnetic EAS deactivation pulse waveform measured with a Rogowski coil (see Equation (3)). The voltage is normalized as the amplitude varies with the location of the measurement. (c) Schematic representation of the deactivator coil with dimensions. All distances were chosen according to measurements taken at a cash register in a retail shop.

Here, 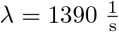, *t*_0_ = 1 ms, *f*_0_ = 1.7 kHz, *φ* = *π* and *H* are the decay rate, delay, frequency, phase angle and Heaviside step function, respectively. Both polarities were evaluated to assess whether polarity influences the results and to ensure that the outcomes are independent of the applied polarity.

Dosimetric calculations were carried out with the simulation platform Sim4Life Version 9.2 (ZurichMedTech AG, Zurich, Switzerland) to study the induced electric fields in the tissues of the heads resulting from the magnetic fields generated by the deactivator coil. For the purpose of this study, the human anatomical models MARTIN (full body) and MIDA (head and neck) were used [24], [25]. They are highly detailed models comprising 86 structures for MARTIN and 115 structures for MIDA at high resolution. The dielectric properties of the virtual model were assigned according to the IT’IS LF database Ver. 5.0 [36]. The electric field simulations were performed with a sinusoidal current at 1700 Hz and with a peak amplitude of 2.1 kA flowing through the current loop.

The induced three-dimensional electric vector field was calculated with 0.5 mm resolution. Fig. 1(d-i) illustrates the in situ electric field distribution in the heads of the MARTIN and MIDA models for one slice.

Table I presents the electric-field percentiles in the gray matter for both anatomical models.

**TABLE I.** Percentiles of E-field in the gray matter 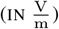.

| Percentile | MARTIN | MIDA |
| --- | --- | --- |
| 99 | 0.096 | 0.086 |
| 99.9 | 0.13 | 0.11 |
| 100 | 0.37 | 0.35 |

#### B. Human neuron simulations

##### 1) Cortical models

As in Aberra et al. (2018, 2020), Fig. 1(j) shows 25 neuron models of the juvenile rat somatosensory cortex microcircuit adapted to reflect the geometric and biophysical properties of human neurons [21], [23], [37], [38]. These multi-compartmental, conductance-based models were modified by the algorithm proposed by Aberra et al. (2018) to incorporate myelination and to adjust geometric parameters [21]. One cell type for each cortical layer was chosen based on their low activation threshold and relative abundance:

1. layer 1 burst non-accommodating neurogliaform cell with a dense axonal arbor
2. layer 2/3 continuous adapting pyramidal cell
3. layer 4 continuous accommodating large basket cell
4. layer 5 continuous adapting thick-tufted pyramidal cell
5. layer 6 continuous adapting tufted pyramidal cell

For each cell type, five statistically varied clones were generated by jittering branch angles and section lengths [23]. Each neuron model consists of multiple morphological sections shown in different colors and characterized by a unique set of membrane mechanisms (Table VI). Each neuronal section is subdivided into segments (or compartments) of equal length to allow numerical simulation. The segment length is determined based on the electrotonic length, ensuring a balance between simulation accuracy and computational cost. A discretization analysis was conducted to ensure that the number of segments was sufficient: excitation threshold error was less than 1% relative to a simulation with seven times as many segments, for both the uniform electric field condition and ICMS condition evaluated on cell model 7.

To couple the electromagnetic field with the cortical cell models, each neuron was placed within the head model in their respective cortical layer with its somatodendritic axis oriented perpendicular to the cortical surface. This was achieved by interpolating between gray and white matter surface meshes using the iterative closest point (ICP) algorithm at normalized depths, which were derived from [37], [39]. The neurons were positioned at the 10 voxel locations closest to the maximum electric field magnitude within the gray matter. Since the wavelength of the electric field is much larger than the size of the brain, the quasi-static regime applies, allowing the spatial distribution and temporal waveform to become separable.

##### 2) Spatial coupling component

The NEURON simulation environment [40] includes a built-in extracellular membrane mechanism that enables the simulation of neuronal responses to externally applied electric fields by specifying the extracellular potential for each compartment. However, this requires converting the electric field into a scalar potential, which is not directly possible using Equation (1), as the presence of the time-varying magnetic vector potential renders the electric field non-conservative. To approximate the effect of this non-conservative field, the local electric field vector is integrated along the path of each compartment, resulting in a scalar quantity referred to as the *quasipotential*. This quasipotential is well defined for a neuron because its morphology does not contain closed loops. Aberra et al. updated their code with a quasipotential integration method [21], [37]:

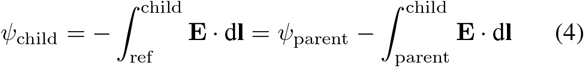

Here, the integral is evaluated from the parent compartment to the considered child compartment, with **dl** representing the displacement vector along the neuron’s morphology. The soma is arbitrarily chosen as the reference compartment and is assigned a quasipotential of zero.

In order to compute the quasipotential, the electric field in each compartment is first linearly interpolated from the voxelized head model, which has a resolution of 0.5 mm. The quasipotential is then calculated at each section by traversing the neuronal tree structure and integrating the field along the neuronal path.

##### 3) Temporal coupling component

The temporal component of the stimulation consists of a finite-duration sinusoidal waveform, where both the simulation timestep and the total duration are frequency-dependent, so the sinusoidal variations are accurately captured and at least one full period is covered:

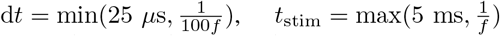

As shown in the Appendix, a minimum of 100 samples per period is required to maintain the relative error below 2%. For very low frequencies, the timestep is set to the default value of 25 *µ*s. The minimum stimulation duration is set to 5 ms to ensure that the resulting threshold value remains close to the rheobase, as demonstrated in the Appendix. For low-frequency stimuli, the duration is extended to match one full period.

##### 4) Neuronal activation threshold

A titration algorithm is used to estimate the excitation threshold for 10 frequencies evenly spaced on a logarithmic scale between 100 Hz and 100 kHz [41]. In this study, the neuron is considered excited when the somatic membrane potential becomes positive. This algorithm adaptively adjusts the amplitude of the sinusoidal waveform in each iteration, achieving a precision of 0.01%. The final titration factor is then multiplied by the 99^*th*^ percentile root-mean-square (RMS) electric field in the gray matter, yielding a threshold value that can be directly compared with the ICNIRP basic restrictions and the IEEE ICES dosimetric reference limits.

#### C. Statistical analysis

Excitation thresholds are analyzed across exposure directions, cell types, subjects and neuron placement locations. Both intra- and inter-group effect sizes are considered. Variation in thresholds is examined for each parameter, and median values are compared to investigate potential selectivity with respect to cell type, subject and exposure direction. A total of 165000 excitation threshold values were obtained, providing a sufficiently large dataset to be used as sample data for a paired statistical hypothesis test comparing different groups. The null hypothesis (H_0_) assumes that there is no systematic difference between the paired populations, while the alternative hypothesis (H_1_) states a nonzero shift. The direction of the difference is inferred from the sign of the median of the paired differences.

## III. Results

Thresholds were computed for 25 neuronal models, consisting of 5 clones per cell type in 6 orthogonal exposure orientations for 11 subjects at 10 locations and 10 stimulation frequencies. First, variations in thresholds were analyzed and statistical tests were conducted to assess significant differences between parameters. Subsequently, these thresholds are compared with the current ICES and ICNIRP guidelines.

### A. Cell models

For each cell type, 3300 (5 clones per type x 6 B-field orientations x 11 subjects x 10 locations) excitation thresholds were computed for each frequency. The median and interquartile ranges as a function of frequency for various cell types are shown in Fig. 3(a). The thresholds for the L4 large basket cell vary the least, where the maximum is only 7.2 times the minimum at 100 Hz and decreases to 3 at 100 kHz. The thresholds for the L2/3 pyramidal cells is one of the lowest based on the median and vary the most with a ratio of 30.6 at 100 Hz and 4.5 at 100 kHz. A decrease in variability with increasing frequency is observable for all cell types. The excitation thresholds of the different cell types are compared by calculating pairwise p-values using a paired difference test on the corresponding threshold distributions. The Wilcoxon signed-rank test was chosen because the threshold is not normally distributed for each combination [42]. Threshold normality test were performed for every cell and frequency, Shapiro-Wilk: p-values ranged from *p <* 10^−44^ to *p <* 10^−10^, Anderson-Darling: the AD statistic was minimally 5 times higher than the critical values for all cases [43], [44]. Fig. 3(c) presents the results of the pairwise comparisons between cell types, showing both the significance of a systematic shift and the direction of that shift for all simulated frequencies. A green color indicates that the median ratio is larger than 1 (that the first mentioned cell has a higher median threshold than the second one), whereas red indicates the opposite.

**Fig. 3.**
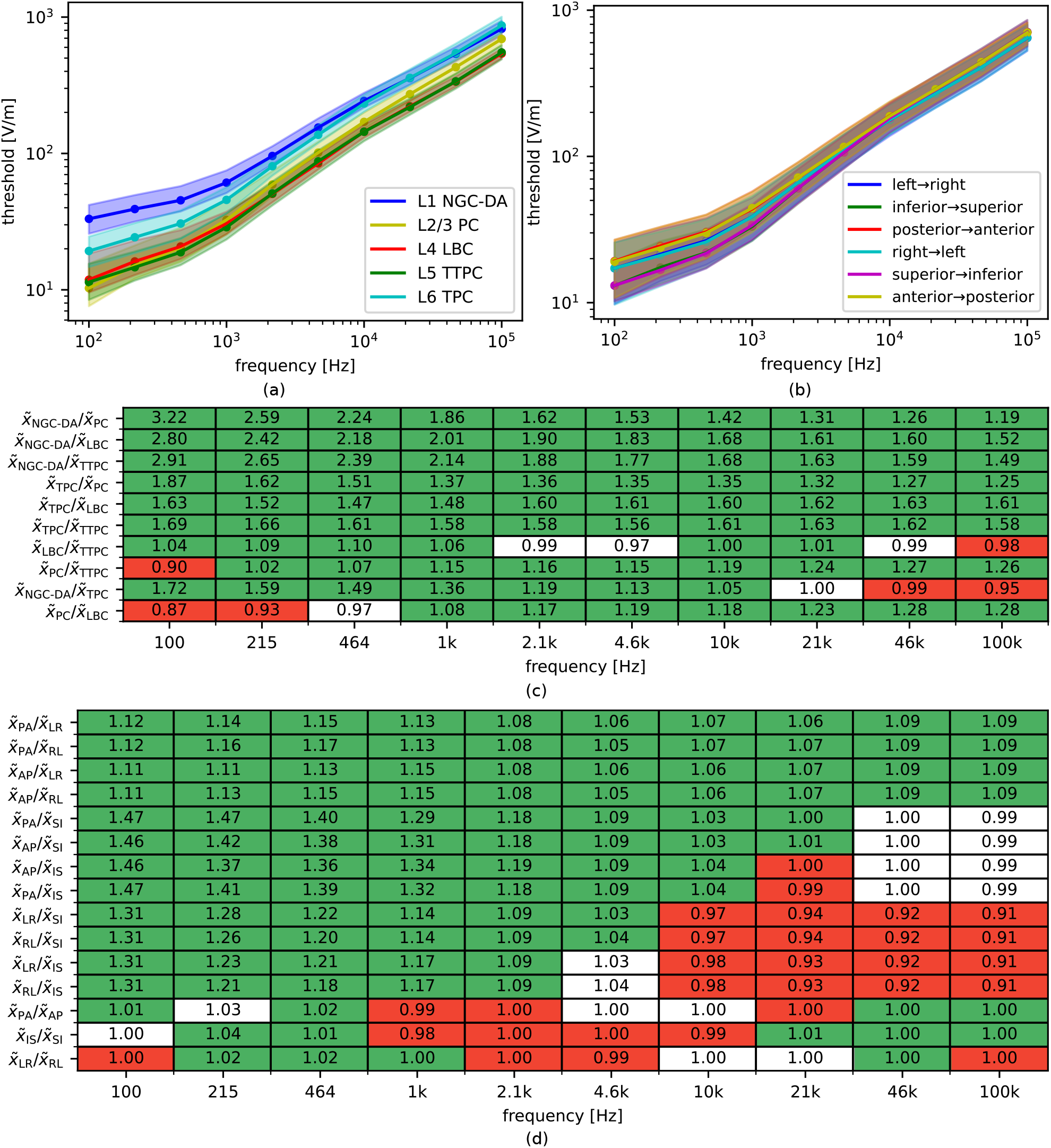
(a) Median RMS threshold values and interquartile ranges for all clones of each cell type across 11 subjects, 6 exposure directions and 10 locations. Dots represent the simulated values. (b) Median RMS threshold values and interquartile ranges for each exposure direction across 25 cell models, 11 subjects and 10 locations. Dots represent the simulated values. (c) Median ratios between cell types. A green or red color indicates if the null hypothesis is rejected (*α* = 0.05) using a pairwise Wilcoxon test. Green indicates that the ratio is larger than one and red indicates that the opposite is true. (d) Median ratios between exposure directions. A green or red color indicates if the null hypothesis is rejected using a pairwise Wilcoxon test. Green indicates that the ratio is larger than one and red indicates that the opposite is true. **Abbreviations**. NGC-DA: neurogliaform cells with a dense axonal arbor, PC: pyramidal cell, LBC: large basket cell, TTPC: thick-tufted pyramidal cell with an early bifurcating apical tuft, TPC L4: tufted pyramidal cell with its dendritic tuft terminating in L4. bNAC: burst non-accommodating, cADpyr: continuous adapting pyramidal, cAC: continuous accommodating, AP: anterior-posterior, LR: left-right, IS: inferior-superior, PA: posterior-anterior, RF: right-left, SI: superior-inferior.

A significance level of 0.05 was adopted as the criterion for rejecting the null hypothesis. Both L1 neurogliaform cells and L6 tufted pyramidal cells consistently exhibit higher values than L2/3 pyramidal cells, L4 large basket cells, and L5 thick-tufted pyramidal cells. The thick-tufted pyramidal cell is almost always significantly lower than the pyramidal cell and the large basket cell. Neurogliaform values remain significantly higher than those of tufted pyramidal cells up to 21 kHz, after which the trend reverses. A similar pattern is observed between the pyramidal cells and the large basket cells around 464 Hz.

### B. Exposure direction

The median threshold and interquartile range for each exposure direction are depicted in Fig. 3(b), based on 2750 values for each frequency. The lateral-medial (right to left) direction exhibits the highest variability across all frequencies, decreasing from 85.9 at 100 Hz to 9.2 at 100 kHz.

Fig. 3(d) indicates that exposure in the posterior-anterior orientation produces significantly higher values compared to the lateral-medial orientation for all frequencies. It also shows that posterior-anterior exposure has higher threshold values than the superior-inferior orientation for all frequencies, except at frequencies higher than 20 kHz. For low frequencies, the lateral-medial orientation exhibits higher values than superior-inferior, whereas the opposite trend is observed at frequencies higher than 5 kHz. The posterior-anterior median is 1.47 times higher than the lowest median at 100 Hz and decreases to 1.09 at higher frequencies. The effect of the direction on excitability is much smaller than that of orientation, as the resulting ratios fall between 0.98 and 1.04.

### C. Subjects

Fig. 4(a) shows the median of 1500 thresholds for all 11 subjects, together with the interquartile ranges of all subjects. Subject 4 has the lowest thresholds for 9 out of 10 frequencies and consistently displays the largest spread. For this subject at 100 Hz, the maximum excitation threshold is 57.6 times higher than the minimum 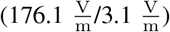. This variation reduces to a factor 23.7 at 1 kHz, 15.3 at 10 kHz, and 9.3 at 100 kHz. The ratio between the median threshold of the most easily excitable subject and that of the least easily excitable subject ranges from 1.18 to 1.48 from 100 Hz to 100 kHz.

**Fig. 4.**
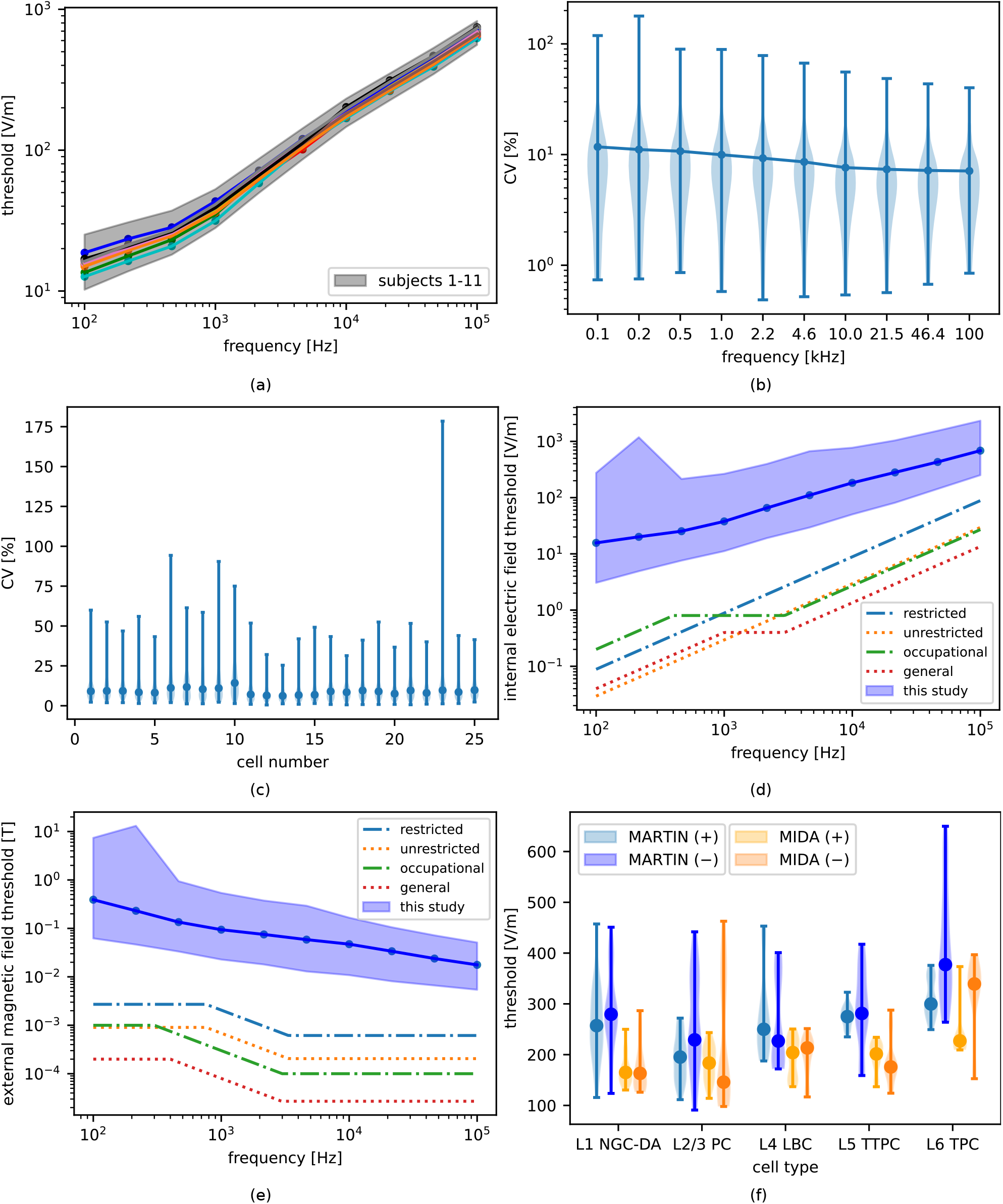
(a) Median RMS threshold values for each subject across 25 cell models, 6 directions and 10 locations. Dots represent the simulated values. The interquartile range across all subjects is shown in gray. (b) Coefficient of variation (CV) across 10 spatial positions for 10 frequencies. Dots indicate median values and violin plots represent the distribution over 25 cells, 6 field directions and 11 subjects. (c) Coefficient of variation across 10 spatial locations for each neuron model. Dots indicate mean values and violin plots represent the distribution over 10 frequencies, 6 field directions and 11 subjects. (d) Minimum, median and maximum RMS internal electric field threshold values for uniform B-field exposure are compared with the current standards and guidelines (ICES restricted and unrestricted environment DRLs and ICNIRP BRs for occupational and general public exposure). (e) Minimum, median and maximum RMS external magnetic field threshold values for uniform B-field exposure are compared with the current standards and guidelines (ICES restricted and unrestricted environment ERLs and ICNIRP RLs for occupational and general public exposure). (f) Comparison of median peak threshold values and corresponding violin plots across 10 spatial locations for the MIDA (head-and-neck model) and MARTIN (full-body model) models. Results are shown for the polarity used in Fig. 2(b) (+) and the reverse polarity case (−) [24], [25].

To quantify the impact of head-model variability on excitation thresholds, a one-way analysis of variance (ANOVA) and a variance decomposition analysis were performed. For all frequencies, the ANOVA yielded p *<* 10^−15^, indicating highly significant differences between subjects. The within-group variability, the between-group variability and the F-statistic increase for higher frequencies. The F-statistic increases from 13.2 at 100 Hz to 64.5 at 100 kHz, showing that high-frequency thresholds differ more strongly between individuals than low-frequency thresholds. To compare between-subject and within-subject variability, the intraclass correlation coefficient (ICC) was computed for each frequency. The ICC increases from 0.008 at 100 Hz to 0.04 at 100 kHz, indicating that although inter-subject variability becomes larger with frequency, it remains small relative to the overall variability. It should be noted that although the relative inter-subject variability ratio decreases, the absolute inter-subject variance continues to increase. Subjects do differ significantly, but the magnitude of their differences is almost negligible compared to the huge within-subject variability.

### D. Neuron placement variability

In addition to the cell type, the subject head model and the exposure direction, the cell positioning introduces variability in the thresholds. The coefficient of variation (CV) across the 10 locations is calculated for all parameter combinations and is shown for each frequency in Fig. 4(b). The median CV remains below 12% for all frequencies, and both the median and maximum decrease with increasing frequency. The largest outliers originate from cell numbers 6, 9 and 23, with cell 23 being the only cell having outliers exceeding 100% at frequencies below 200 Hz. Fig. 4(c) depicts the coefficient of variation (CV) across locations for the 25 considered cell types. The largest cells (number 6 and 9) exhibit the highest CV values, whereas the smallest cell types (number 1-3) show lower threshold. This suggests that location-dependent differences may be amplified for larger cell models (Pearson correlation coefficient: 0.44, p-value: 2.7%), although cell size is not the only contributing factor.

### E. Comparison with guidelines and standards

The worst-case threshold across all cells, directions and subjects is compared with previously reported stimulation thresholds and existing guidelines. Comparison with guideline thresholds requires taking into account the applied reduction factors presented in Table IV. These safety factors are included to address uncertainties in guideline development by accounting for biological and dosimetric limitations, as well as variability in response thresholds across the human population.

In Table II the lowest threshold value of all cells, subjects and directions is divided by both the ICNIRP guidelines and the IEEE standards. These values represent how conservative the current guidelines are compared to the simulated thresholds. The external magnetic field corresponding to this lowest threshold was computed and compared with the guideline limits in Table III. For direct comparison with the hazard threshold, the ratios in Table II are divided by the corresponding reduction factors from Table IV, yielding the threshold ratios shown in Table V. The worst-case value occurs at 100 kHz, where the excitation threshold is only 2.8 times higher than the reported ICES restricted environment guideline, resulting in a threshold ratio of 0.9 after division by the safety factor of 3.

**TABLE II.** Worst-case margin of safety for different frequencies (in Hz). **A****bbreviations**. DRL: dosimetric reference limit (ICES). BR: basic restriction (ICNIRP).

| Guideline/Standard | 100 | 215 | 464 | 1k | 2.1k | 4.6k | 10k | 21k | 46k | 100k |
| --- | --- | --- | --- | --- | --- | --- | --- | --- | --- | --- |
| DRL restricted | 34.5 | 25.7 | 18.5 | 12.6 | 10.0 | 7.2 | 5.7 | 4.2 | 3.5 | 2.8 |
| DRL unrestricted | 103.8 | 77.3 | 55.5 | 37.9 | 30.0 | 21.5 | 17.1 | 12.8 | 10.4 | 8.5 |
| BR occupational | 15.3 | 11.4 | 9.5 | 14.0 | 23.8 | 23.5 | 18.6 | 13.9 | 11.4 | 9.3 |
| BR general | 76.4 | 56.9 | 40.9 | 27.9 | 47.6 | 46.9 | 37.2 | 27.8 | 22.8 | 18.6 |

**TABLE III.** Worst-case margin of safety for different frequencies (in Hz). **A****bbreviations**. ERL: exposure reference level (ICES). RL: reference level (ICNIRP).

| Guideline/Standard | 100 | 215 | 464 | 1k | 2.1k | 4.6k | 10k | 21k | 46k | 100k |
| --- | --- | --- | --- | --- | --- | --- | --- | --- | --- | --- |
| ERL restricted | 23.0 | 17.1 | 12.3 | 11.0 | 18.8 | 21.0 | 17.6 | 13.2 | 10.8 | 8.8 |
| ERL unrestricted | 68.9 | 51.3 | 36.8 | 33.1 | 56.5 | 63.0 | 52.9 | 39.5 | 32.4 | 26.4 |
| RL occupational | 62.3 | 46.4 | 51.5 | 75.8 | 129.3 | 129.1 | 108.4 | 81.0 | 66.3 | 54.1 |
| RL general | 311.4 | 232.1 | 193.3 | 284.4 | 484.9 | 478.0 | 401.4 | 300.1 | 245.7 | 200.5 |

**TABLE IV.** Safety factors.

| Guideline/Standard | 100 Hz - 400 Hz | 400 Hz - 1 kHz | 1 kHz - 100 kHz |
| --- | --- | --- | --- |
| ICES restricted | 3 | 3 | 3 |
| ICES unrestricted | 9 | 9 | 9 |
| ICNIRP occupational | - | 5 | 5 |
| ICNIRP general | - | - | 10 |

**TABLE V.** Worst-case threshold ratio for different frequencies (in Hz). A**bbreviations**. DRL: dosimetric reference limit (ICES). BR: basic restriction (ICNIRP).

| Guideline/Standard | 100 | 215 | 464 | 1k | 2.1k | 4.6k | 10k | 21k | 46k | 100k |
| --- | --- | --- | --- | --- | --- | --- | --- | --- | --- | --- |
| DRL restricted | 11.5 | 8.6 | 6.2 | 4.2 | 3.3 | 2.4 | 1.9 | 1.4 | 1.2 | 0.9 |
| DRL unrestricted | 11.5 | 8.6 | 6.2 | 4.2 | 3.3 | 2.4 | 1.9 | 1.4 | 1.2 | 0.9 |
| BR occupational | - | - | 1.9 | 2.8 | 4.8 | 4.7 | 3.7 | 2.8 | 2.3 | 1.9 |
| BR general | - | - | - | 2.8 | 4.8 | 4.7 | 3.7 | 2.8 | 2.3 | 1.9 |

Fig. 4(d) shows all calculated internal electric field thresholds compared with the recommendations for both occupational and public exposure cases. As expected, the threshold values increase with frequency, consistent with the the guidelines. At 215 Hz, the threshold distribution is more skewed than at other frequencies, which results from simulations with reversed polarities in exposure directions. The difference between the lowest thresholds and the standards narrows as the frequency rises.

Fig. 4(e) compares all calculated external magnetic field thresholds with the guideline recommendations. The thresholds exhibit a decreasing trend consistent with the guidelines, and the worst-case threshold differs from the guideline values by approximately the same factor across all frequencies.

### F. Acousto-magnetic electronic article surveillance system

The threshold was determined for both the head-and-neck model MIDA and whole-body model MARTIN for 25 cell models at 10 different cortical locations considering both polarities for the configuration of Fig. 2 [24], [25]. Fig. 4(f) shows the threshold distributions and median values for each cell type. The MARTIN threshold distribution demonstrates a higher central tendency relative to that of MIDA. Considering the minimum values, L2/3 PCs are the most excitable and L6 TPCs the least for both anatomical models. For the L6 TPCs, the median threshold displays the strongest polarity dependence. The worst-case value is a threshold of 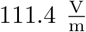 for L2/3 PC in the MARTIN model using the polarity shown in Fig. 2(b). For the reverse polarity, the minimum threshold is 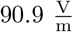

These derived thresholds cannot be compared directly with the standards, as done in Section III-E, because the waveform in Fig. 2(b) is a damped sine wave and therefore nonsinusoidal. The energy in the frequency spectrum of a damped sine is distributed over many nearby frequencies. Its spectrum has a Lorentzian shape centered at *f*_0_ with a bandwidth proportional to the decay rate *λ*.

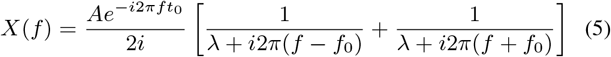

Both a peak-based compliance method and a Fourier-component summation method are described in the recommendations [1], [2] For the peak-based method, the threshold must be divided by 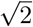 to allow comparison with the permissible levels at *f*_0_ (the highest permissible level at 1.7 kHz is 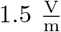). Using this approach, the worst-case margin of safety is 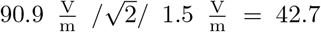 when compared with the ICES restricted standard. When accounting for the safety factors, the worst-case threshold ratios are 14.2 (ICES) and 16.1 (ICNIRP). For the spectral summation method, each Fourier component is compared with its corresponding limit, and added to the sum of fractional exposures. The frequency spectrum in Equation (5) is divided by the guideline limits and integrated up to 10 MHz. Multiplying this integral by the worst-case threshold of 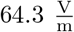 yields margins between 26.0 (ICES restricted) and 100.5 (ICNIRP general).

## IV. Discussion

The results of this study indicate that the current established standards are sufficiently conservative for CNS protection and help bridge the gap between induced electric fields and physiological response thresholds. All 165000 uniform magneticfield exposure scenarios were found to comply with each of the four evaluated guidelines. Threshold variability was maximized by incorporating multiple neuron models across different cortical layers, various head models, diverse exposure directions, and spatial placement. In addition, thresholds were also computed for two anatomical models subjected to a realistic, non-uniform exposure configuration simulated from a realistic EAS coil model, thereby adding additional anatomical and field-distribution variability to the analysis.

### A. Sources of variability

The biophysical cortical cell models were selected following the approach in [21], choosing for each cortical layer the cell type exhibiting the lowest activation threshold. When comparing the five cell types, clear differences in activation threshold are observed, with a strong dependence on field frequency. Moreover, ranking the cell types by their median thresholds across human head models, spatial placements and exposure conditions yields a different ordering than ranking them by their minimum thresholds at certain frequencies. In Fig. 3(c), the cell types are compared based on their threshold distribution, but this mainly reflects their central tendency. For safety assessment relative to the guidelines, the worst-case (minimum) thresholds should instead be considered. Across the considered frequencies, electromagnetic safety is primarily determined by the L23 PC, L4 LBC and L5 TTPC models, as the remaining cell types exhibit substantially higher median and minimum thresholds. Therefore, if only the safety margin with respect to the guidelines is of interest for a particular frequency, simulating only the cell type with the lowest threshold at that frequency is sufficient.

Head models were obtained from 11 healthy young adults of both genders. Although individual head models exhibit statistically significant differences in excitation thresholds (F-statistics exceeding 10), this between-subject variation remains small compared to the overall variability. For all frequencies, the intraclass correlation (ICC) remained below 0.04, indicating that inter-subject variability is much lower than within-subject variability. This result is not entirely unexpected, as all subjects fall within a relatively narrow age range of 25 to 37 years, and therefore do not account for children or elderly people, who may exhibit markedly different anatomical properties and tissue conductivities. To extend the age range, a child model (MARTIN) was also evaluated in a realistic exposure scenario. No consistent trends were observed when comparing thresholds across gender or age. For each frequency, a Mann–Whitney U test was performed to compare the two genders: the p-values ranged from 0.78 to 10^−12^ with increasing frequency, while the U-statistic remained nearly constant. Although some p-values indicated statistical significance, the effect size derived from the U-statistic 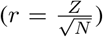 was very small, between -0.055 and -0.0022 (with r the rank-biserial correlation coefficient, Z the standardized U-statistic, and N the total sample size). For age, the pearson correlation coefficient between age and excitation threshold ranged from -0.077 to -0.036 (with p *<* 10^−6^), indicating no meaningful association between age and excitability within this age range.

Six exposure configurations were examined by orienting the uniform magnetic field along the three orthogonal anatomical axes, each applied with forward and reversed polarity. A clear distinction is observed between the three exposure orientations, although the differences remain relatively small: the maximum ratio of median thresholds between two directions is 1.47 in Fig. 3(d). The posterior-anterior orientation is the least excitable across nearly all frequencies. At low frequencies, the superior-inferior direction produces the lowest median thresholds, whereas at higher frequencies the lateral-medial orientation becomes the most excitable according to the median. Across all frequencies, the minimum threshold is consistently lowest for the lateral-medial direction, making this orientation the most relevant for safety assessment simulations. The difference between the forward and reverse polarities is relatively small compared to the orientational differences and depends strongly on frequency, as shown in the last 3 rows in 3(d). Nevertheless, exposure directions that deviate from the three principal axes could lead to threshold values that are more difficult to predict. Both guidelines considered in this study assume a polarization condition in which the magnetic field vector is aligned perpendicular to the long axis of a body, as this configuration yields optimal coupling between the tissue and the incident field and therefore represents a worst-case scenario. However, this specific alignment rarely reflects typical exposure conditions. Neurons were placed according to cortical lamination and oriented along columnar geometry, resulting in varying orientations relative to the applied field and thereby capturing a broad and realistic range of alignment scenarios.

Neurons were placed within the head model at 10 distinct cortical locations, with their somatodendritic axes oriented perpendicular to the local cortical surface, resulting in unique spatial positions and corresponding orientations. The median coefficient of variation across these 10 locations remains below 12%, indicating that spatial location contributes only moderately to variability and that excitability is generally consistent across the cortex layers. The outliers exceeding 90% occur at the two lowest frequencies, while outliers above 70% appear in the lower half of the simulated frequency range. These outliers primarily originate from locations that are noticeably more difficult to excite compared to the others. This is shown in Fig. 4(d), the low-frequency thresholds are asymmetrically skewed toward higher values. A comparison of Fig. 4(b) and Fig. 4(c) demonstrates that the asymmetric peak around 215 Hz accounts for the elevated CV outliers.

Two highly detailed anatomical models were subjected to a strong magnetic deactivation pulse. Utilizing the peak-based method, the safety margins are approximately three times higher for the AM-EAS thresholds compared to the worst-case safety margins for uniform exposure at 1.7 kHz. This observation is consistent with the guidelines, which consider spatially uniform fields to represent the worst-case scenario. When applying the spectral summation method, the worst-case margin is 31.5. Because this method assumes in-phase addition of all spectral components, it yields overly conservative estimates for pulsed waveforms. The thresholds of the 29-month old male model (MARTIN) are clearly higher than those of the adult model (MIDA), which is unexpected, as children generally exhibit lower excitation thresholds due to their higher intrinsic excitability [45]–[48]. A possible explanation is that whole-body models such as MARTIN may require a stronger external stimulus to reach activation because a larger portion of the induced current could leak into the body, effectively acting as a current sink, whereas head models such as MIDA confine currents to the head. As a result, head-only models may artificially isolate the brain and consequently overestimate the cortical electric fields.

### B. Comparison with prior studies

Figure 5 compares the results of this study with findings from two other numerical studies and two experimental publications [13], [18], [49], [50]. When a study employed a specific neuronal model, it is indicated in the figure with a citation. Gomez-Tames et al. (2025) compared thresholds for five membrane channel dynamics using axon trajectories in three head models: Hodgkin-Huxley (HH) based on a squid axon, Frankenhaeuser-Huxley (FH) based on a frog node, Chiu-Ritchie-Rogart-Stagg-Sweeney (CRRSS) based one a rabbit node, Schwarz-Eikhof (SE) based on a rat node, and Schwarz-Reid-Bostock (SRB) based on a human nerve [51]–[55]. Additionally, Soyka et al. (2025) also considered two double-cable models [56]. The results of Soyka et al. (2025) [18], which used three different straight axon models under a homogeneous aligned electric field, were plotted as originally reported [19], [57], [58]. For Gomez-Tames et al. (2025) [13], which employed the SENN model [15], [59], only the median thresholds are shown because the reported thresholds span more than two orders of magnitude. In contrast, the present study reports the minimum, median, and maximum threshold values for completeness. In addition to previous computational findings, two experimental data points were included from in-vitro neurons derived from the cerebral cortex of a rat embryo and fetus [49], [50]. Overall, the results of this study are consistent with those reported previously, although they exhibit slightly higher excitation thresholds and span two orders of magnitude. This spread highlights the importance of using validated neuron models when evaluating excitation thresholds relevant to LF EMF exposure assessment.

**Fig. 5.**
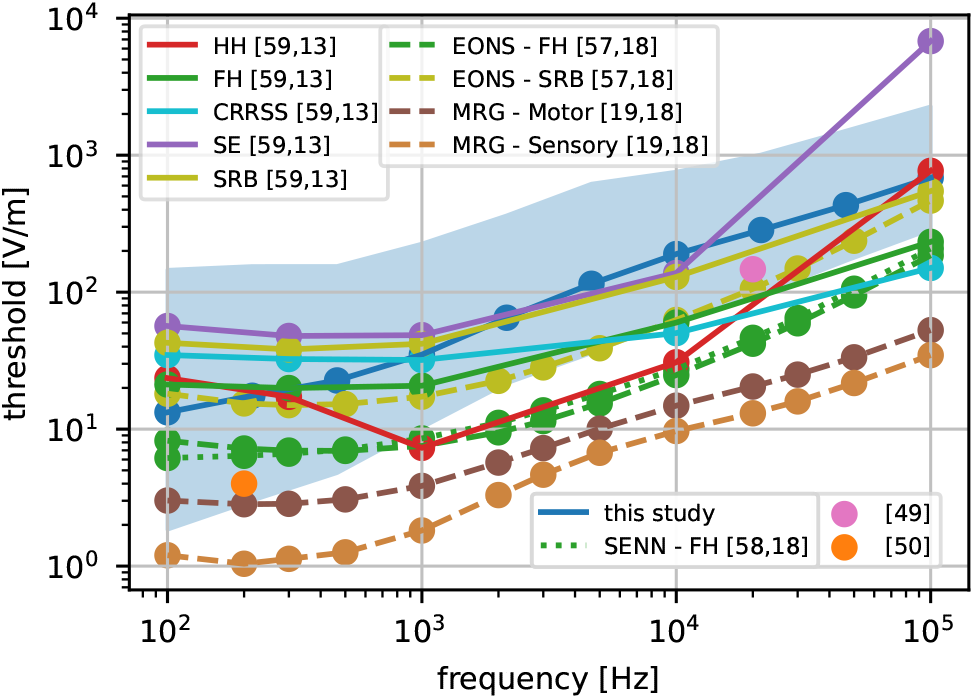
Comparison of internal electric field thresholds of this study with Gomez-Tames et al. (2025), Soyka et al. (2025), Saito et al. (2018) and Ikehata et al. (2020) [13], [18], [49], [50]. Minimum, median and maximum thresholds are plotted for this study and median thresholds for Gomez-Tames et al. (2025). **Abbreviations**. HH: Hodgkin-Huxley, FH: Frankenhaeuser-Huxley, CRRSS: Chiu-Ritchie-Rogart-Stagg-Sweeney, SE: Schwarz-Eikhof, SRB: Schwarz-Reid-Bostock.

### C. Strengths and limitations

This study evaluates 175000 parameter combinations by varying exposure conditions, cell types, anatomical models, neuron locations and stimulation frequencies. For the uniform exposure scenario, the electric field was computed using both a FEM solver and an SPFD solver to ensure robust simulation results.

The following limitations should be considered when interpreting the results of this study. First, this paper measures excitation thresholds assuming neurons are in their resting state. Thus, it does not take into account subthreshold effects, which modulate neural activity without eliciting an action potential [60], [61]. Subthreshold neuromodulation is identified as a gap in current safety assessments by IEEE ICES and ICNIRP [16], [17]. Quantifying cortical subthreshold neuromodulation to LF EMF exposure in the context of safety guidelines will be the focus of future work, where the models presented in the current paper will be repurposed.

Second, as mentioned in the research agenda of IEEE ICES, the induced electric field measured from voxelized human head models and numerical methods is vulnerable to the staircasing error [16], [62]. This was mitigated by the 99^th^ percentile method, which is recommended by the ICNIRP guidelines [2]. Third, corticofugal axons, which are highly myelinated and project to subcortical areas, were not considered in this study. Hence, the full range of cortical neuron morphology is not represented [63]. To place our findings in context and account for these large output neurons, we compare our results to previous studies on corticofugal axons in Fig. 5 [12], [13].

Furthermore, this study considers a uniform magnetic field exposure scenario as a general baseline condition, which may not fully represent real-world situations. To mitigate this limitation, EAS simulations were also included as a more realistic exposure scenario, providing a closer representation of typical LF EMF exposure conditions. However, the MARTIN and MIDA phantoms do not include anisotropic tissue conductivities, which may be relevant in white matter, and do not incorporate age-dependent conductivity variations.

Finally, for the two AM-EAS scenarios, the same cortical cell models were used for both the child and adult phantoms. This assumption is not realistic, as children typically have smaller neurons, different ion-channel distributions, and a higher synaptic density. These developmental differences may explain why the child model did not exhibit lower excitability than the adult model.

Future work can extend this methodology to additional LF exposure scenarios, such as WPT and induction heating.

## V. Conclusion

This study provides a comprehensive evaluation of LF EMF induced neuronal activation using morphologically realistic cortical nerve cell models incorporated into anatomically detailed human head models. By combining voxel-based dosimetry with finite-element and finite-difference simulations, we analyzed 165000 uniform magnetic field exposure scenarios. In addition, we investigated a realistic non-uniform exposure case involving a child near and AM-EAS deactivator.

Across all scenarios, considerable variability in excitation threshold was observed, predominantly driven by cell type, field orientation and frequency. Inter-subject anatomical variability contributed significantly less than within-subject variability. L2/3 and L5 pyramidal cells and L4 large basket cells display the lowest excitation thresholds, making them the most relevant for safety assessments. Frequency strongly influenced both threshold magnitude and variability, with higher frequencies yielding reduced variability and higher thresholds.

Despite the broad range of exposure conditions tested and the inclusion of highly excitable neuronal morphologies, all excitation thresholds remained above the recommended internal field limits specified by both IEEE ICES and ICNIRP. When applying the specified safety factors, an exception emerged at 100 kHz, where the threshold fell slightly below the hazard level, with the lowest simulated excitation threshold reaching 0.9 times the hazard threshold. Nevertheless, thresholds at all other frequencies remained on the protective side of the limit. Results from the AM-EAS deactivator scenario similarly demonstrated conservative safety margins using both the MIDA and MARTIN anatomical models.

Overall, these findings confirm that existing LF EMF exposure guidelines remain conservative, even when compared to detailed morphological neuron models and extensive anatomical and field based variability. The use of morphologically realistic neurons improves the understanding of electromagnetic excitability in the human cortex and strengthens the evidence supporting current exposure recommendations. The present results support the view that existing LF EMF limits remain conservative with respect to the modeled excitation thresholds considered in this study.

## Acknowledgments

The computational resources (Stevin Supercomputer Infrastructure) and services used in this work were provided by the VSC (Flemish Supercomputer Center), funded by Ghent University, FWO and the Flemish Government – department EWI.

The authors would like to acknowledge the valuable discussions with IEEE ICES Technical Committee 95 Subcommittee 6 during this work.

This work is supported by the Special Research Fund BOF 01Z00824 and BOF basic financing bof/baf/4y/2024/01/080.

## Appendix

### A1. Discretization analysis

A discretization analysis was performed for several simulation parameters to ensure that the resulting threshold values were sufficiently accurate. The timestep and total simulation duration were swept over a wide range by placing cell number 7 at the spatial peak of the E-field in a 29-year-old male model with a left to right exposure. Fig. 6(a) shows the relative error as a function of the number of samples per period: using 100 samples per period results in a maximum error of 2% (for frequencies below 400 Hz, the time step equals the default value of 25 *µ*s). In Fig. 6(b), the relative error is plotted as a function of the simulated number of periods. A minimum duration of 5 ms was imposed (which corresponds to the period of a 200 Hz sine wave), as higher frequencies require a larger number of cycles to achieve stable threshold estimates. The number of segments was not further increased, as Fig. 6(c) shows that additional segmentation results in only negligible changes. Both a uniform electric field case and an intracortical microstimulation case were included to cover the full range of spatial stimulation conditions. A comparison between linear and cubic interpolation of the electric fields simulated with the FEM-method to the neuronal compartment locations is shown in Fig. 6(d) using the same parameters as in Fig. 6(a,b) for 10 different locations. The interpolation errors 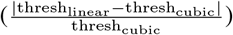 remain acceptable (maximal smaller than 16.2%, mean ¡ 7 & across the considered frequency range), justifying the use of linear interpolation.

**Fig. 6.**
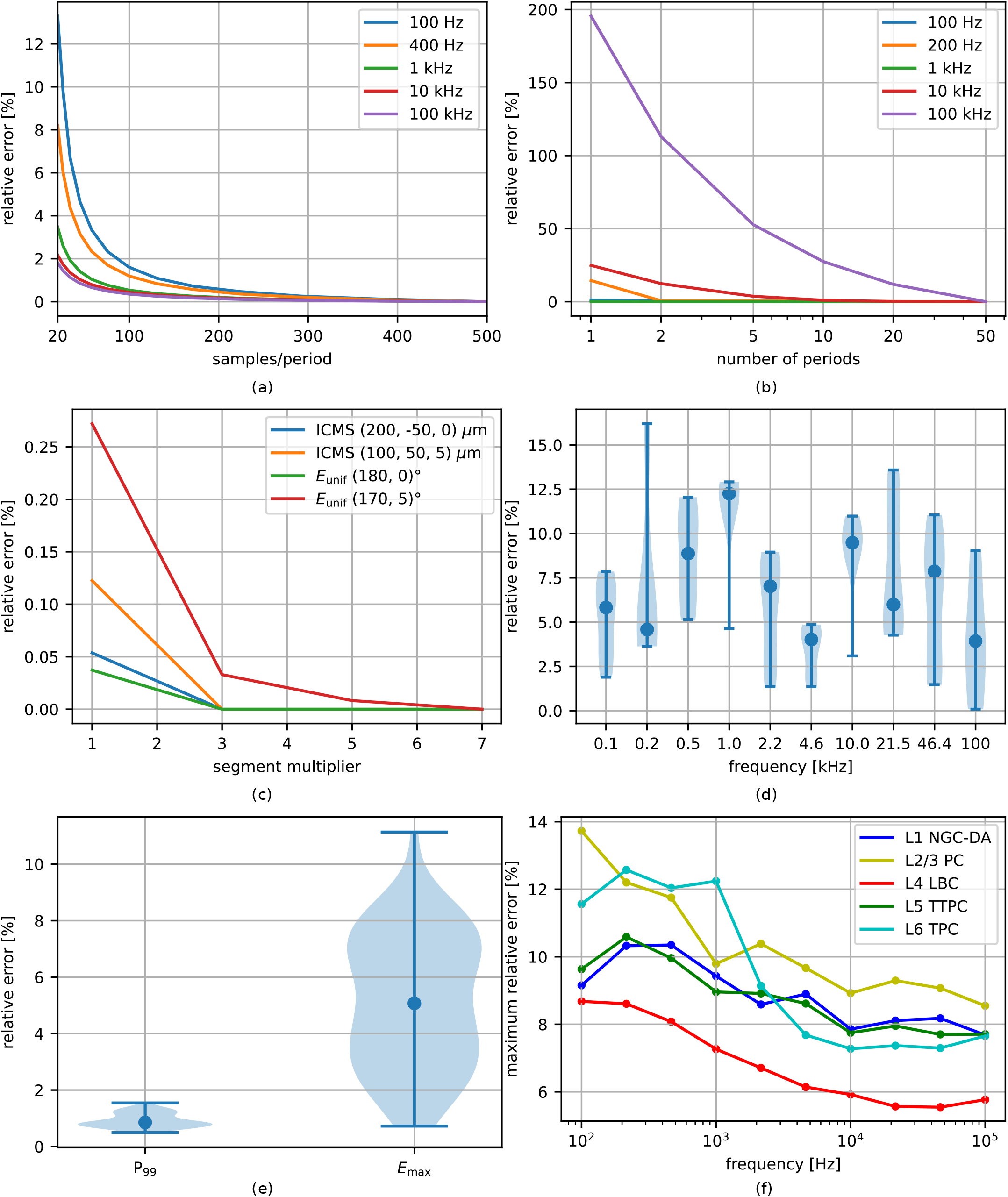
(a) Relative threshold error as a function of the number of time samples per period for various frequencies. (b) Relative threshold error as a function of the number of simulated periods. (c) Relative threshold error as a function of the segment multiplier for various conditions. (d) Relative threshold error between linear and cubic interpolation for every frequency. Dots indicate mean values and violin plots represent the distribution over 25 cell models, 6 field directions and 11 subjects. (e) Relative error of the maximum E-field and the 99th percentile of FEM- and SPFD-derived electric fields. (f) Maximum relative threshold error between the FEM and SPFD method.

All 165000 simulated thresholds for the uniform magnetic field case were computed using 25 neuronal models exposed to 6 orthogonal orientations for 11 subjects at 10 locations for which both FEM- and SPFD-derived electric fields were evaluated to verify the robustness of the neuron simulations. In Fig. 6(e), both the maximum electric field and 99th percentile are compared across all head models. The maximum-field error 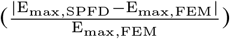 remains below 12%, and the 99th percentile error 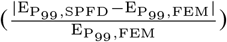 stays below 2%, supporting the use of P_99_ to reduce staircasing artifacts. Fig. 6(f) shows that the maximal relative error 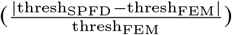 does not exceed 14% among all cell types, head models and exposure orientations.

### A2. Membrane mechanisms

The membrane mechanisms inserted in the different sections of all cell models are shown in Table VI.

**TABLE VI.** Inserted MechanismType objects in each section type for all 25 neuron models.

| Section type | Mechanisms (inhibitory cells 1–5, 11–15) | Mechanisms (excitatory cells 6–10, 16–25) |
| --- | --- | --- |
| Soma | pas, extracellular, CaDynamics_E2, Ca_LVAst, Ca, Im, K_Pst, K_Tst, NaTs2_t, Nap_Et2, SK_E2, SKv3_1, xtra | pas, extracellular, CaDynamics_E2, Ca_HVA, Ca_LVAst, Ih, NaTs2_t, SK_E2, SKv3_1, xtra |
| Axon | pas, extracellular, CaDynamics_E2, Ca_LVAst, Ca, Im, K_Pst, K_Tst, NaTa_t, Nap_Et2, SK_E2, SKv3_1, xtra | pas, extracellular, CaDynamics_E2, Ca_HVA, Ca_LVAst, K_Pst, K_Tst, NaTa_t, Nap_Et2, SK_E2, SKv3_1, xtra |
| Basal dendrite | pas, extracellular, Ih, Im, K_Pst, K_Tst, NaTs2_t, Nap_Et2, SKv3_1, xtra | pas, extracellular, Ih, xtra |
| Apical dendrite | / | pas, extracellular, Ih, Im, NaTs2_t, SKv3_1, xtra |
| Node of Ranvier | pas, extracellular, K_Pst, K_Tst, NaTa_t, Nap_Et2, SKv3_1, xtra | pas, extracellular, K_Pst, K_Tst, NaTa_t, Nap_Et2, SKv3_1, xtra |
| Myelin | pas, extracellular, xtra | pas, extracellular, xtra |
| Unmyelin | pas, extracellular, K_Pst, K_Tst, NaTa_t, Nap_Et2, SKv3_1, xtra | pas, extracellular, K_Pst, K_Tst, NaTa_t, Nap_Et2, SKv3_1, xtra |

### A3. Key Resources Table

The key resources used in this study are shown in Table VII.

**TABLE VII.**
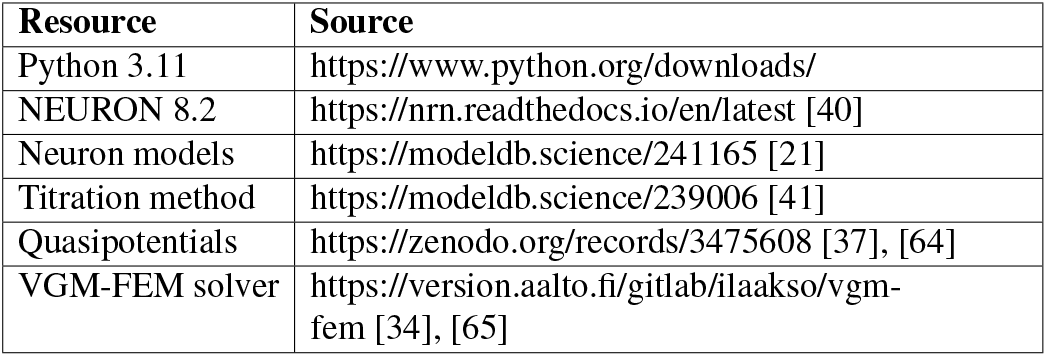
Software and algorithms.

| Resource | Source |
| --- | --- |
| Python 3.11 | https://www.python.org/downloads/ |
| NEURON 8.2 | https://nrn.readthedocs.io/en/latest [40] |
| Neuron models | https://modeldb.science/241165 [21] |
| Titration method | https://modeldb.science/239006 [41] |
| Quasipotentials | https://zenodo.org/records/3475608 [37], [64] |
| VGM-FEM solver | https://version.aalto.fi/gitlab/ilaakso/vgm-fem [34], [65] |

## Notes

https://github.com/jgazq/ENAT26

### Competing Interest Statement

The authors have declared no competing interest.

## References

[1] IEEE, “IEEE standard for safety levels with respect to human exposure to electric, magnetic, and electromagnetic fields, 0 hz to 300 ghz,” pp. 1–312, 2019.

[2] ICNIRP, “Guidelines for limiting exposure to time-varying electric and magnetic fields (1 hz to 100 khz),” pp. 818–836, 2010.

[3] J. P. Reilly, “Neuroelectric mechanisms applied to low frequency electric and magnetic field exposure guidelines—part i: Sinusoidal waveforms,” Health physics, vol. 83, no. 3, pp. 341–355, 2002.

[4] D. Shi, C. Zhu, R. Lu, S. Mao, and Y. Qi, “Intermediate frequency magnetic field generated by a wireless power transmission device does not cause genotoxicity in vitro,” Bioelectromagnetics, vol. 35, no. 7, pp. 512–518, 2014.

[5] I. Laakso and A. Hirata, “Evaluation of the induced electric field and compliance procedure for a wireless power transfer system in an electrical vehicle,” Physics in Medicine & Biology, vol. 58, no. 21, p. 7583, 2013.

[6] S. Aerts, C. Calderon, B. Valič, M. Maslanyj, D. Addison, T. Mee, C. Goiceanu, L. Verloock, M. Van den Bossche, P. Gajšek et al., “Measurements of intermediate-frequency electric and magnetic fields in households,” Environmental Research, vol. 154, pp. 160–170, 2017.

[7] B. Floderus, C. Stenlund, and F. Carlgren, “Occupational exposures to high frequency electromagnetic fields in the intermediate range (> 300 hz–10 mhz),” Bioelectromagnetics, vol. 23, no. 8, pp. 568–577, 2002.

[8] A. Hirata, Y. Diao, T. Onishi, K. Sasaki, S. Ahn, D. Colombi, V. De Santis, I. Laakso, L. Giaccone, W. Joseph, E. A. Rashed, W. Kainz, and J. Chen, “Assessment of human exposure to electromagnetic fields: Review and future directions,” IEEE Transactions on Electromagnetic Compatibility, vol. 63, no. 5, pp. 1619–1630, 2021.

[9] J. P. Reilly, Applied bioelectricity: from electrical stimulation to electropathology. Springer Science & Business Media, 2012.

[10] H.-J. Lee, H. Jin, Y. H. Ahn, N. Kim, J. K. Pack, H.-D. Choi, and Y.-S. Lee, “Effects of intermediate frequency electromagnetic fields: a review of animal studies,” International Journal of Radiation Biology, vol. 99, no. 2, pp. 166–182, 2023.

[11] C. d. A. Lellis, M. A. M. Herbas, and L. J. d. Silva, “Psychiatric disorders after deep brain stimulation of the subthalamic nucleus in parkinson’s disease: a systematic review,” einstein (São Paulo), vol. 22, p. eRW0182, 2024.

[12] J. Gomez-Tames, T. Tarnaud, K. Miwa, A. Hirata, T. Van de Steene, L. Martens, E. Tanghe, and W. Joseph, “Brain cortical stimulation thresholds to different magnetic field sources exposures at intermediate frequencies,” IEEE Transactions on Electromagnetic Compatibility, vol. 61, no. 6, pp. 1944–1952, 2019.

[13] J. Gomez-Tames, T. Tarnaud, W. Joseph, and E. Tanghe, “Numerical study on human brain cortical electrostimulation assessment during uniform magnetic field exposure at intermediate frequencies,” IEEE Access, 2025.

[14] Y. Diao, W. Joseph, D. Poljak, L. Giaccone, S. Kodera, I. Laakso, K. Yamazaki, K. Li, K. Sasaki, E. Tanghe et al., “Recent advances and future perspective in computational bioelectromagnetics for exposure assessments,” Bioelectromagnetics, vol. 46, no. 3, p. e70002, 2025.

[15] J. P. Reilly, V. T. Freeman, and W. D. Larkin, “Sensory effects of transient electrical stimulation - evaluation with a neuroelectric model,” IEEE Transactions on Biomedical Engineering, vol. BME-32, no. 12, pp. 1001–1011, 1985.

[16] J. P. Reilly and A. Hirata, “Low-frequency electrical dosimetry: research agenda of the ieee international committee on electromagnetic safety,” Physics in Medicine & Biology, vol. 61, no. 12, p. R138, 2016.

[17] G. Ziegelberger, C. Marino, R. Croft, M. Feychting, A. Green, A. Hirata, G. D’Inzeo, S. Miller, T. Okuno, Z. Sienkiewicz et al., “Gaps in knowledge relevant to the “guidelines for limiting exposure to time-varying electric and magnetic fields (1 hz-100 khz),” Health physics, vol. 118, no. 5, pp. 533–542, 2020.

[18] F. Soyka, T. Tarnaud, C. Alteköster, R. Schoeters, T. Plovie, W. Joseph, and E. Tanghe, “Action potential threshold variability for different electrostimulation models and its potential impact on occupational exposure limit values,” Bioelectromagnetics, vol. 46, no. 1, p. e22529, 2025.

[19] J. L. Gaines, K. E. Finn, J. P. Slopsema, L. A. Heyboer, and K. H. Polasek, “A model of motor and sensory axon activation in the median nerve using surface electrical stimulation,” Journal of Computational Neuroscience, vol. 45, no. 1, pp. 29–43, 2018. [Online]. Available: 10.1007/s10827-018-0689-5

[20] M. Soldati, M. Mikkonen, I. Laakso, T. Murakami, Y. Ugawa, and A. Hirata, “A multi-scale computational approach based on tms experiments for the assessment of electro-stimulation thresholds of the brain at intermediate frequencies,” Physics in Medicine & Biology, vol. 63, no. 22, p. 225006, 2018.

[21] A. S. Aberra, A. V. Peterchev, and W. M. Grill, “Biophysically realistic neuron models for simulation of cortical stimulation,” Journal of neural engineering, vol. 15, no. 6, p. 066023, 2018.

[22] H. R. Siebner, K. Funke, A. S. Aberra, A. Antal, S. Bestmann, R. Chen, J. Classen, M. Davare, V. Di Lazzaro, P. T. Fox et al., “Transcranial magnetic stimulation of the brain: What is stimulated?–a consensus and critical position paper,” Clinical neurophysiology, vol. 140, pp. 59–97, 2022.

[23] H. Markram, E. Muller, S. Ramaswamy, M. W. Reimann, M. Abdellah, C. A. Sanchez, A. Ailamaki, L. Alonso-Nanclares, N. Antille, S. Arsever et al., “Reconstruction and simulation of neocortical microcircuitry,” Cell, vol. 163, no. 2, pp. 456–492, 2015.

[24] M. I. Iacono, E. Neufeld, E. Akinnagbe, K. Bower, J. Wolf, I. Vogiatzis Oikonomidis, D. Sharma, B. Lloyd, B. J. Wilm, M. Wyss, K. P. Pruessmann, A. Jakab, N. Makris, E. D. Cohen, N. Kuster, W. Kainz, and L. M. Angelone, “Mida: A multimodal imaging-based detailed anatomical model of the human head and neck,” PLOS ONE, vol. 10, no. 4, pp. 1–35, 04 2015. [Online]. Available: 10.1371/journal.pone.0124126

[25] H. Jeong, G. Ntolkeras, M. Alhilani, S. R. Atefi, L. Zöllei, K. Fujimoto, A. Pourvaziri, M. H. Lev, P. E. Grant, and G. Bonmassar, “Development, validation, and pilot mri safety study of a high-resolution, open source, whole body pediatric numerical simulation model,” PLOS ONE, vol. 16, no. 1, pp. 1–31, 01 2021. [Online]. Available: 10.1371/journal.pone.0241682

[26] I. Laakso, S. Tanaka, S. Koyama, V. De Santis, and A. Hirata, “Inter-subject variability in electric fields of motor cortical tdcs,” Brain Stimulation, vol. 8, no. 5, pp. 906–913, 2015. [Online]. Available: https://www.sciencedirect.com/science/article/pii/S1935861X15009419

[27] A. M. Dale, B. Fischl, and M. I. Sereno, “Cortical surface-based analysis: I. segmentation and surface reconstruction,” NeuroImage, vol. 9, no. 2, pp. 179–194, 1999. [Online]. Available: https://www.sciencedirect.com/science/article/pii/S1053811998903950

[28] B. Fischl, M. I. Sereno, and A. M. Dale, “Cortical surface-based analysis: Ii: Inflation, flattening, and a surface-based coordinate system,” NeuroImage, vol. 9, no. 2, pp. 195–207, 1999. [Online]. Available: https://www.sciencedirect.com/science/article/pii/S1053811998903962

[29] P. Dimbylow, “Development of the female voxel phantom, naomi, and its application to calculations of induced current densities and electric fields from applied low frequency magnetic and electric fields,” Physics in Medicine & Biology, vol. 50, no. 6, p. 1047, feb 2005. [Online]. Available: 10.1088/0031-9155/50/6/002

[30] S. Gabriel, R. W. Lau, and C. Gabriel, “The dielectric properties of biological tissues: Ii. measurements in the frequency range 10 hz to 20 ghz,” Physics in Medicine & Biology, vol. 41, no. 11, p. 2251, nov 1996. [Online]. Available: 10.1088/0031-9155/41/11/002

[31] C. A. Bossetti, M. J. Birdno, and W. M. Grill, “Analysis of the quasi-static approximation for calculating potentials generated by neural stimulation,” Journal of neural engineering, vol. 5, no. 1, p. 44, 2007.

[32] G. Gaugain, L. Quéguiner, M. Bikson, R. Sauleau, M. Zhadobov, J. Modolo, and D. Nikolayev, “Quasi-static approximation error of electric field analysis for transcranial current stimulation,” Journal of Neural Engineering, vol. 20, no. 1, p. 016027, 2023.

[33] A. Hirata, F. Ito, and I. Laakso, “Confirmation of quasi-static approximation in sar evaluation for a wireless power transfer system,” Physics in Medicine & Biology, vol. 58, no. 17, p. N241, 2013.

[34] I. Laakso and A. Hirata, “Fast multigrid-based computation of the induced electric field for transcranial magnetic stimulation,” Physics in Medicine & Biology, vol. 57, no. 23, p. 7753, nov 2012. [Online]. Available: 10.1088/0031-9155/57/23/7753

[35] T. Dawson and M. Stuchly, “High-resolution organ dosimetry for human exposure to low-frequency magnetic fields,” IEEE Transactions on Magnetics, vol. 34, no. 3, pp. 708–718, 1998.

[36] C. Baumgartner, P. A. Hasgall, F. Di Gennaro, E. Neufeld, B. Lloyd, M. C. Gosselin, D. Payne, A. Klingenböck, and N. Kuster, “IT’IS Database for Thermal and Electromagnetic Parameters of Biological Tissues,” Aug. 2025. [Online]. Available: https://itis.swiss/database

[37] A. S. Aberra, B. Wang, W. M. Grill, and A. V. Peterchev, “Simulation of transcranial magnetic stimulation in head model with morphologically-realistic cortical neurons,” Brain Stimulation, vol. 13, no. 1, pp. 175–189, 2020. [Online]. Available: https://www.sciencedirect.com/science/article/pii/S1935861X19304097

[38] S. Ramaswamy, J.-D. Courcol, M. Abdellah, S. R. Adaszewski, N. Antille, S. Arsever, G. Atenekeng, A. Bilgili, Y. Brukau, A. Chalimourda et al., “The neocortical microcircuit collaboration portal: a resource for rat somatosensory cortex,” Frontiers in neural circuits, vol. 9, p. 44, 2015.

[39] P. Besl and N. D. McKay, “A method for registration of 3-d shapes,” IEEE Transactions on Pattern Analysis and Machine Intelligence, vol. 14, no. 2, pp. 239–256, 1992.

[40] M. L. Hines and N. T. Carnevale, “The neuron simulation environment,” Neural computation, vol. 9, no. 6, pp. 1179–1209, 1997.

[41] J. P. Reilly, “Survey of numerical electrostimulation models,” Physics in Medicine & Biology, vol. 61, no. 12, p. 4346, 2016.

[42] F. Wilcoxon, “Individual comparisons by ranking methods,” Biometrics Bulletin, vol. 1, no. 6, pp. 80–83, 1945. [Online]. Available: http://www.jstor.org/stable/3001968

[43] S. S. Shapiro and M. B. Wilk, “An analysis of variance test for normality (complete samples)†,” Biometrika, vol. 52, no. 3-4, pp. 591–611, 12 1965. [Online]. Available: 10.1093/biomet/52.3-4.591

[44] T. W. Anderson and D. A. Darling, “Asymptotic theory of certain “Goodness of Fit” criteria based on stochastic processes,” The Annals of Mathematical Statistics, vol. 23, no. 2, pp. 193–212, Jun. 1952.

[45] Y. Ben-Ari, “Excitatory actions of gaba during development: the nature of the nurture,” Nature Reviews Neuroscience, vol. 3, no. 9, pp. 728–739, 2002. [Online]. Available: 10.1038/nrn920

[46] D. Warm, J. Schroer, and A. Sinning, “Gabaergic interneurons in early brain development: Conducting and orchestrated by cortical network activity,” Frontiers in Molecular Neuroscience, vol. Volume 14 - 2021, 2022. [Online]. Available: https://www.frontiersin.org/journals/molecular-neuroscience/articles/10.3389/fnmol.2021.807969

[47] K. Kirmse and C. Zhang, “Principles of gabaergic signaling in developing cortical network dynamics,” Cell Reports, vol. 38, no. 13, p. 110568, 2022. [Online]. Available: 10.1016/j.celrep.2022.110568

[48] K. Cohen Kadosh, B. Krause, A. J. King, J. Near, and R. Cohen Kadosh, “Linking gaba and glutamate levels to cognitive skill acquisition during development,” Human Brain Mapping, vol. 36, no. 11, pp. 4334–4345, 2015.

[49] A. Saito, T. Terai, K. Makino, M. Takahashi, S. Yoshie, M. Ikehata, Y. Jimbo, K. Wada, Y. Suzuki, and S. Nakasono, “Real-time detection of stimulus response in cultured neurons by high-intensity intermediate-frequency magnetic field exposure,” Integrative Biology, vol. 10, no. 8, pp. 442–449, 07 2018. [Online]. Available: 10.1039/c8ib00097b

[50] M. Ikehata, Y. Suzuki, A. Saito, and S. Yoshie, “Evaluation of stimulation threshold in nerve cells by exposure to time-varying magnetic field in in vitro,” Quarterly Report of RTRI, vol. 61, no. 4, pp. 297–302, 2020.

[51] A. L. Hodgkin and A. F. Huxley, “A quantitative description of membrane current and its application to conduction and excitation in nerve,” The Journal of Physiology, vol. 117, no. 4, pp. 500–544, 1952. [Online]. Available: https://physoc.onlinelibrary.wiley.com/doi/abs/10.1113/jphysiol.1952.sp004764

[52] B. Frankenhaeuser and A. F. Huxley, “The action potential in the myelinated nerve fibre of xenopus laevis as computed on the basis of voltage clamp data,” The Journal of Physiology, vol. 171, no. 2, pp. 302–315, 1964. [Online]. Available: https://physoc.onlinelibrary.wiley.com/doi/abs/10.1113/jphysiol.1964.sp007378

[53] S. Y. Chiu, J. M. Ritchie, R. B. Rogart, and D. Stagg, “A quantitative description of membrane currents in rabbit myelinated nerve.” The Journal of Physiology, vol. 292, no. 1, pp. 149–166, 1979. [Online]. Available: https://physoc.onlinelibrary.wiley.com/doi/abs/10.1113/jphysiol.1979.sp012843

[54] J. R. Schwarz and G. Eikhof, “Na currents and action potentials in rat myelinated nerve fibres at 20 and 37° c,” Pflügers Archiv, vol. 409, no. 6, pp. 569–577, Aug. 1987. [Online]. Available: 10.1007/BF00584655

[55] J. R. Schwarz, G. Reid, and H. Bostock, “Action potentials and membrane currents in the human node of ranvier,” Pflügers Archiv, vol. 430, no. 2, pp. 283–292, Jun. 1995. [Online]. Available: 10.1007/BF00374660

[56] C. C. McIntyre, A. G. Richardson, and W. M. Grill, “Modeling the excitability of mammalian nerve fibers: Influence of afterpotentials on the recovery cycle,” Journal of Neurophysiology, vol. 87, no. 2, pp. 995–1006, 2002, pMID: 11826063. [Online]. Available: 10.1152/jn.00353.2001

[57] T. Tarnaud, F. Sokya, R. Schoeters, T. Plovie, W. Joseph, L. Martens, and E. Tanghe, “Eons: Evaluation of non-sinusoidal magnetic fields for electromagnetic safety to intermediate frequencies,” in The 1st Annual Meeting of BioEM (BioEM 2022), 2022, pp. 320–325.

[58] J. P. Reilly and A. M. Diamant, Electrostimulation: theory, applications, and computational model. Artech House, 2011.

[59] J. P. Reilly, H. Antoni, M. A. Chilbert, and J. D. Sweeney, “From electrical stimulation to electropathology,” Applied Bioelectricity, vol. 1, pp. 1–563, Aug. 1998.

[60] A. Šarolić, M. Perić, D. Kulić, and D. Sapunar, “Neuromodulation by electromagnetic field exposure in the intermediate frequency range,” in 2023 24th International Conference on Applied Electromagnetics and Communications (ICECOM), 2023, pp. 1–4.

[61] A. S. Aberra, R. Wang, W. M. Grill, and A. V. Peterchev, “Multi-scale model of axonal and dendritic polarization by transcranial direct current stimulation in realistic head geometry,” Brain stimulation, vol. 16, no. 6, pp. 1776–1791, 2023.

[62] I. Laakso and A. Hirata, “Reducing the staircasing error in computational dosimetry of low-frequency electromagnetic fields,” Physics in medicine & biology, vol. 57, no. 4, p. N25, 2012.

[63] S. Tomasi, R. Caminiti, and G. M. Innocenti, “Areal differences in diameter and length of corticofugal projections,” Cerebral cortex, vol. 22, no. 6, pp. 1463–1472, 2012.

[64] Aman-A, “Aman-a/tmssim aberra2019: First release,” Oct. 2019. [Online]. Available: 10.5281/zenodo.3475608

[65] L. Ilkka, “Vgm-fem,” Nov. 2023. [Online]. Available: https://version.aalto.fi/gitlab/ilaakso/vgm-fem

